# Design and Synthesis of Potent First-in-Class Histones Acetyltransferase Modulators

**DOI:** 10.1101/2025.03.03.641271

**Authors:** Elisa Zuccarello, Mattia Mori, Elisa Calcagno, Rosa Purgatorio, Yan Feng, Erica Acquarone, Yitshak I. Francis, Mauro Fa’, Agnieszka Staniszewski, Elentina K. Argyrousi, Coleman D. Barry, Hong Zhang, Shi-Xian Deng, Donald W. Landry, Ottavio Arancio, Jole Fiorito

**Affiliations:** Taub Institute for Research on Alzheimer’s Disease and the Aging Brain, Columbia University, New York, NY 10032, United States; Department of Medicine, Columbia University, New York, NY, United States; Department of Biotechnology, Chemistry and Pharmacy, University of Siena, Italy; Department of Pathology & Cell Biology, Columbia University, New York, NY, United States; New York Institute of Technology, Department of Biological and Chemical Sciences, Northern Boulevard, Old Westbury, NY 11568, United States

**Keywords:** Epigenetics, HAT modulators, p300, drug design, molecular docking, metabolic investigation

## Abstract

Epigenetic regulation governs gene expression through histone modifications, particularly the acetylation of lysine residues. These modifications are orchestrated by enzymes such as histone acetyltransferases (HATs) and histone deacetylases (HDACs). Among them, p300 plays a critical role in various disorders such as neurodegenerative conditions (i.e. Alzheimer’s disease, and Alzheimer’s Disease related dementia) and cancer (i.e. glioblastoma and B cell lymphoma). Our aim is to develop therapeutic interventions for these ailments. In this context, we designed and synthesized novel activators of the enzyme. Using an *in vitro* screening approach combined with structural activity relationship analysis, we successfully identified compounds that modulate histone acetylation.

## 1. Introduction

The epigenetic machinery regulates post-translational modifications of histones, which in turn modulate gene expression and protein synthesis. A key modification is the acetylation of lysine residues on histones, a process finely controlled by two classes of enzymes: histone acetyltransferases (HATs) and histone deacetylases (HDACs) with opposite functions towards the acetylation of the substrate (1). HATs transfer acetyl-groups to the ε-amino groups of specific lysines in histones, as well as to other transcription factors (e.g., p53) and other nuclear proteins (e.g., α-tubulin) (2–4). Consequently, HATs modulate gene transcription, nucleosome assembly, and DNA repair. Altered HAT functions lead to several diseases, ranging from neurodegenerative disorders to cancer (4–6).

Four main HAT families contribute to histone acetylation in humans, the p300/CREB binding protein (CBP) family (including members p300 and CBP), the GNAT family (including general control non-repressed 5 protein - GCN5, p300/CBP associated factor - PCAF, HAT1, and ATF2), the MYST family (including Tat-interactive protein-60 - Tip60, MOZ, MORF, and HBO1), and the nuclear receptor coactivator family (including steroid receptor coactivator-1 - SRC-1, ACTR, and TIF-2) (7). There is also an additional group of HATs including various members such as TAF_II_250, TFIIIC, and CLOCK.

The p300/CBP family is of relevance to our studies as the compounds were screened for activity towards p300. The enzyme often has functional redundancy with CBP due to structural similarity, and sequence homology. Recent studies have demonstrated that p300 plays a pivotal role in various cellular processes, including cell cycle control, differentiation, and apoptosis (4). Specifically, p300 is crucial for the transcriptional coactivation of CREB, c-Jun, c-Fos, c-Myb, p53, Stats nuclear receptor, RelA, GATA, and p73 (8). Alterations of p300 enzymatic activity are associated with different human diseases, such as neurodegenerative diseases, cancer, cardiac diseases, diabetes, and inflammatory processes (9, 10). CBP binds to multiple transcription factors, including c-jun, c-myb, MyoD, E2F1, YY1, and members of the steroid hormone receptor superfamily (11). CBP is involved in neurodegenerative diseases, cancer, cardiovascular diseases, asthma, and diabetes (12). Both proteins have nine separate functional domains including CH1, KIX, CH3 (ZZ-TAZ), and NCBD domains (13). These domains are linked by long stretches of residues to the catalytic core of p300 and CBP, which contains the bromodomain (BRD), the CH2 region, and the HAT domain. The HAT domain acetylates histones, and the BRD recognizes acetylated substrates (14, 15). Moreover, BRD and CH2 regions are required for chromatin binding, but how these domains cooperate to read out chromatin modifications is still unknown (13). The BRD domain and CH2 regions seem to be crucial as modulators of HAT activity. In addition, BRD is required in the HAT domain for effective substrate acetylation, and loss of the BRD impairs p300 HAT substrate specificity and transcriptional activity (16, 17). Structural alteration of p300 and CBP, such as CH2 region mutations, have been found to inactivate HAT activity, but the real mechanism is unknown (18). Recent studies have revealed a discontinuous domain on the CH2 region, called plant homeodomain (PHD) type zinc finger motif, interrupted by a small domain, the RING domain. Typically, the PHD domain binds trimethylated histone H3 Lys4 (H3K4me3), unmodified histone-H3 tails, or other modified chromatin regions (19). Delvecchio *et al*. (2013) found that the BRD, PHD, RING, and HAT domains form a compact module in which the RING domain is juxtaposed with the HAT substrate-binding site and the RING domain has an inhibitory function for the HAT domain. Moreover, Delvecchio *et al*. (2013) discovered that mutations at the RING domain led to the upregulation of HAT autoacetylation and p53 substrate acetylation. These discoveries indicate that HAT regulation requires repositioning of the RING domain to facilitate access to an otherwise partially occluded HAT active site. Such mutations of the RING domain cause p300 HAT dysregulation leading to pathogenic conditions. These structural and mechanistic studies on the p300 catalytic core provide a comprehensive understanding of the role of p300 regulation and dysregulation in diseases and how chromatin recognition and modification are associated (13).

Here, we report the discovery and synthesis of new small molecules that modulate the activity of p300 based on structure-activity relationship (SAR) analysis of the CTPB scaffold for HAT activators (20, 21). Overall, the outcomes demonstrate a library of small molecules bearing an *N*-phenylbenzamide structure. The new chemical entities either increase, or decrease, or have no activity towards the acetylation of lysines 18 and 27 of histone 3. Among them, the activator YF2 was characterized for *in vitro* microsomal stability, molecular docking, and metabolic profile.

## 2. Results

### 2.1. Design of new chemical entities that modulate p300 activity

Our SAR studies started with the structures of two anacardic acid analogs, CTPB and CTB. Based on the structures of the analogs, we designed and synthesized a library of small molecules bearing an *N*-phenylbenzamide structure. Eight representative small molecules out of the library are shown in **Figure 1**.

**Figure 1.**
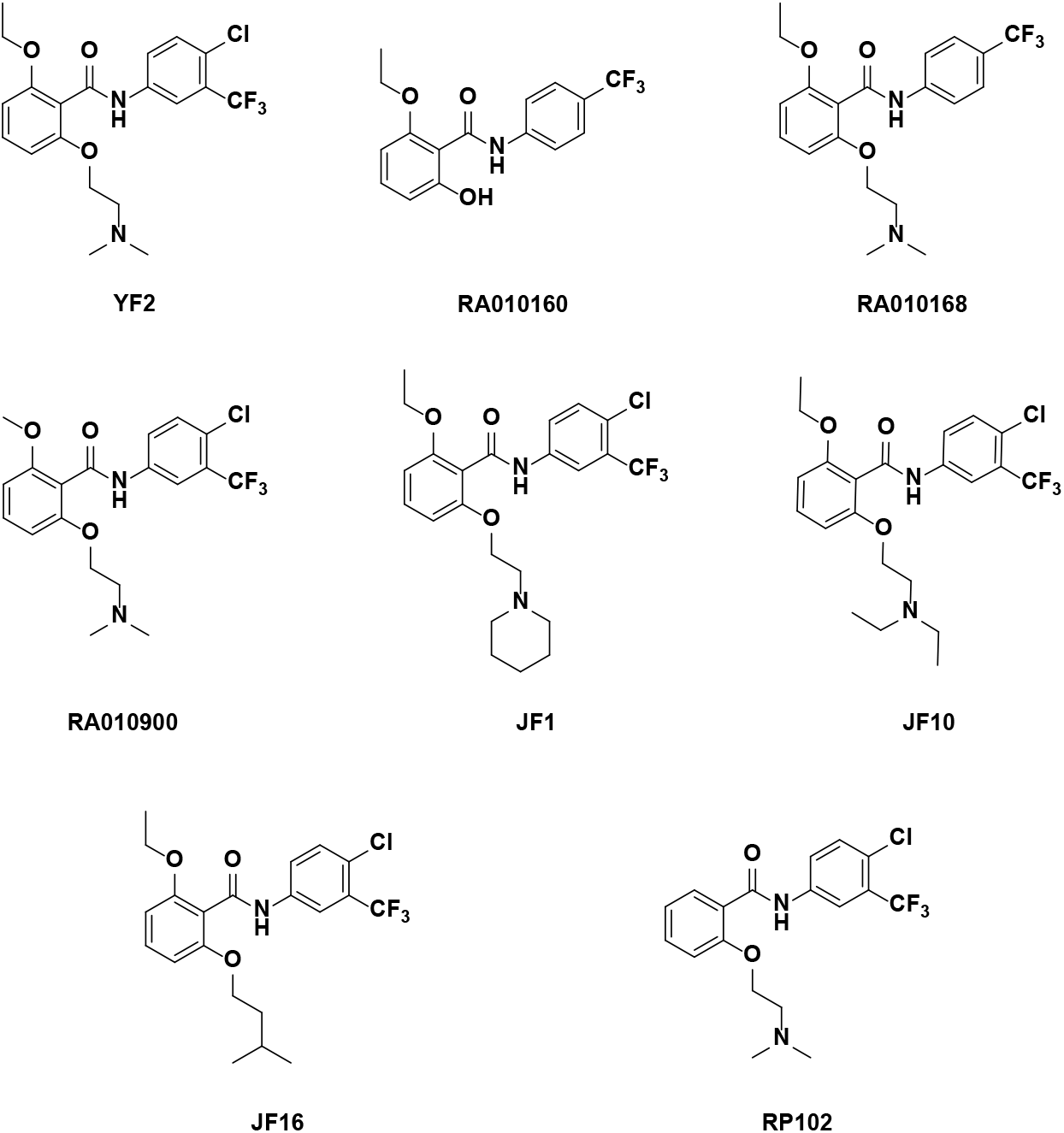
*N*-phenylbenzamide analogs: YF2, RA010160, RA010168, RA010900, JF1, JF10, JF16, and RP102.

### 2.2. Synthesis of the new chemical entities with the *N*-phenylbenzamide scaffold

Following the design of these chemical entities we proceeded to their synthesis. The goal of the synthetic efforts was to obtain novel, potent p300 modulators, each with a high yield and purity. Our synthetic approach started with protection reactions using thionyl chloride alongside acetone and DMAP in anhydrous DME in the presence of commercially sourced 2,6-dihydroxybenzoic acid (**Scheme 1**). This procedure yielded compound **2** in a scalable amount. The next step involved alkylation reactions of compound **2** with various alkyl halides in the presence of Cs_2_CO_3_ and NaI in MEK at room temperature for 24 hours, resulting in compounds **3a-g**. To further functionalize compounds **3a-g**, the hydrolysis of the 1,3-dioxin-4-one moiety was conducted with CsOH in THF at reflux for 72 hours, yielding compounds **4a-g** in great amounts. Additionally, amidation reactions were carried out using EDC with an appropriate aniline or aminopyridine and DMAP in anhydrous CH_2_Cl_2_, overnight at room temperature, producing compounds **5a-g**. The final step involved alkylation reactions in the presence of an appropriate alkyl halide and Cs_2_CO_3_ in 1,4-dioxane at 70°C for 16h or Mitzunobu reactions with an appropriate alcohol, PPh_3_, and DIAD in anhydrous THF, at room temperature for 24 hours, resulting in the new series of p300 ligands.

**Scheme 1.**
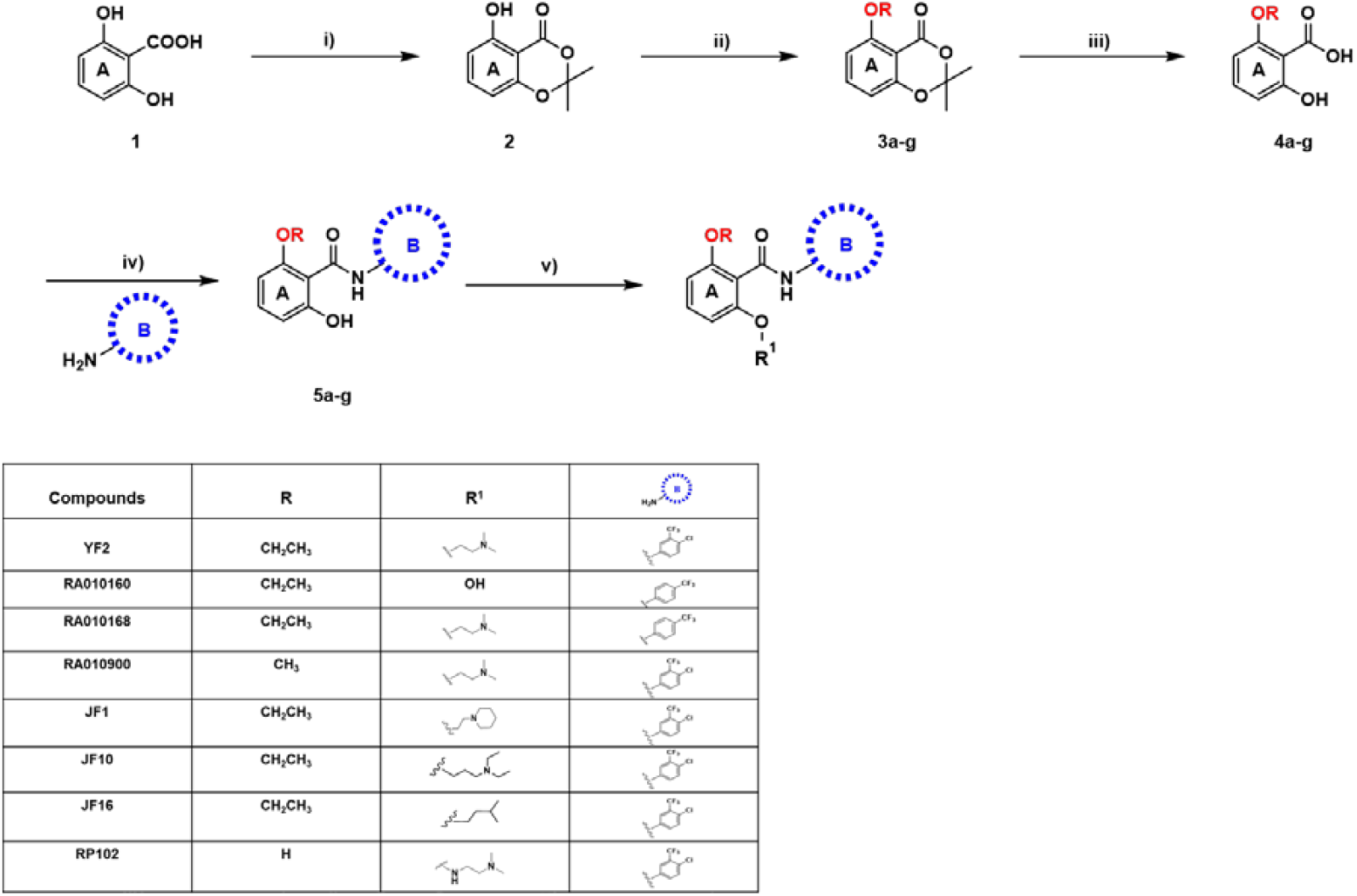
Synthesis of *N*-phenylbenzamide analogs. Reagents and conditions: i) SOCl_2_, Acetone, DMAP, DME, rt, 8 hour; ii) appropriate alkyl halide, Cs_2_CO_3_, NaI, MEK, rt, 24 hours; iii) CsOH, THF, reflux, 72 hours; iv) appropriate aniline, EDC HCl, CH_2_Cl_2_, rt, overnight; v) appropriate alkyl halide, Cs_2_CO_3_, 1,4-dioxane, 70°C, 16 hours, or appropriate ethanolamine, PPh_3_, DIAD, THF, rt, 24 hours.

### 2.3. Enzymatic activity of *N*-phenylbenzamide analogs towards p300

Following the synthesis of the new chemical entities with the *N*-phenylbenzamide scaffold, the compounds were tested for activity through a cell-free assay in which the HAT enzyme, the acetyl acceptor histone 3, the acetyl donor acetyl-CoA, and the compound at the appropriate concentrations were mixed *in vitro*. Enzyme activity was assessed by measuring levels of acetylation of lysine residues through specific antibodies by immunoblot. Using this method, the EC_50_ values of the new chemical entities for acetylation of lysine residues 18 and 27 of histone 3 by p300 have been established (**Table 1**).

**Table 1.** p300 enzymatic activity of analogs *N*-phenylbenzamide derivatives.

| Compound | Activity on Lysine 18<br>(K18) nM ± SEM | Activity on Lysine 27<br>(K27) nM ± SEM |
| --- | --- | --- |
| YF2 | Activator<br>EC<sub>50</sub> 155.01 ± 0.32 | Activator<br>EC<sub>50</sub> 72.54 ± 0.27 |
| RA010160 | Activator<br>EC<sub>50</sub> 145.85 ± 0.65 | Activator<br>EC<sub>50</sub> 532.91 ± 1.07 |
| RA010168 | Activator<br>EC<sub>50</sub> 48.02 ± 1.34 | No Activity |
| RA010900 | Activator<br>EC<sub>50</sub> 116.82 ± 1.31 | Activator<br>EC<sub>50</sub> 43.98 ± 0.66 |
| JF1 | Inhibitor<br>IC<sub>50</sub> 54.79 ± 1.32 | Inhibitor<br>IC<sub>50</sub> 129.06 ± 1.09 |
| JF10 | Inhibitor<br>IC<sub>50</sub> 101.39 ± 0.64 | Inhibitor<br>IC<sub>50</sub> 62.87 ± 0.47 |
| JF16 | Inhibitor<br>IC<sub>50</sub> 11.59 ± 0.36 | Inhibitor<br>IC<sub>50</sub> 951.97 ± 2.59 |
| RP102 | N/A | N/A |

### 2.4 Structure-Activity Relationship Analysis of the *N*-phenylbenzamide analogs

Compounds **RA010160** and **RA010168** contain a mono-substituted ring B, where the CF3 group is in the para position. While this structural modification leads to the discovery of activators, the simultaneous presence of the R^1^ dimethylaminoethyl chain seems to affect the activity toward the acetylation of the Lys K27 residue. The findings on compounds **YF2** and **RA010900**, which differ in the alkyl group −OR linked to ring A, are particularly significant. They show that smaller alkyl chains (**YF2** and **RA010900**) promote activator behavior. These findings suggest that the size and structure of the alkyl group can significantly influence the biological activity of the compound.

The N-phenylbenzamide scaffold was modified on the R^1^ side chain with different substituted tertiary amines, and aliphatic moieties. Additionally, we investigated the impact of increasing steric hindrance of the substituents on the benzene ring on compounds potency. Compound **JF1**, showing the cyclic amine, such as piperidin-1-yl moiety, as the R^1^ group, inhibits p300 for the acetylation of both lysine residues and behave as an inhibitor. In light of these observations, the nature of the alkyl group R^1^ is a key determinant in the compound behavior. When R^1^ is a *N, N*-dimethylaminoethyl side chain (**YF2, RA010168** and **RA010900**) or hydroxyl group (**RA010160**) the molecules maintain their activator nature on one or both lysine. However, when amino-alkyl and alkyl groups are longer and branched (**JF10** and **JF16**), the molecule inverts its behavior, becoming an inhibitor. This finding underscores the significant influence of the alkyl group on the compound’s activity.

Based on these findings, we selected compound **YF2** for further *in vitro* pharmacokinetics investigation and carried out molecular modeling studies to elucidate the binding mode on the catalytic site of the p300 enzyme.

### 2.5. YF2 Molecular Docking Campaign

The possible interaction of the p300 activator **YF2** to the crystallographic structure of the p300 catalytic core was investigated by molecular docking simulations with the GOLD docking program (22). Given the availability of structural information on the interaction between small molecules and the bromodomain of the p300 close homologue CBP (23–26), we focused our interest on the bromodomain binding site of p300. Docking results showed that **YF2** can properly fit the p300 bromodomain binding site. Specifically, the *N*-phenylbenzamide core is docked in a lipophilic pocket composed of Leu1073, Pro1074, Phe1075, Val1079, Leu1084, Gly1085, Ile1086, Tyr1089, Tyr1131, and Val1138. The chlorine atom and trifluoromethyl group are docked in a region composed of Leu1073 and Gln1077, while the protonated basic moiety is docked near the entrance of the binding site in a solvent-exposed region where it establishes an H-bond with the side chain of Asn1132 (**Figure 2**).

**Figure 2.**
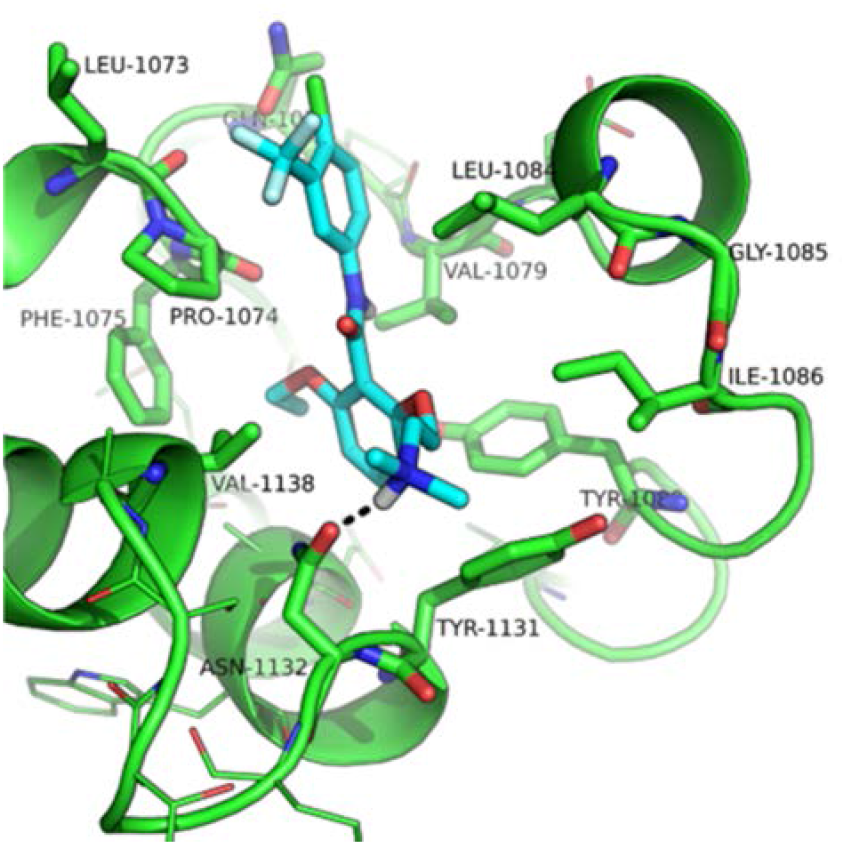
Docking-based binding mode of YF2 within the p300 bromodomain. The crystallographic structure of p300 coded by PDB-ID: 4BHW is shown in green. Residues within 5 Å from the ligands are shown as lines, while those surrounding **YF2** are shown as sticks and are labeled. **YF2** is represented by cyan sticks, non-polar H atoms are omitted; H-bond interactions are highlighted by black dashed lines.

### 2.6. *In vitro* metabolic profiles of compound YF2

Compound **YF2** was tested for its metabolic stability in the presence of mouse and human liver microsomes. The *in vitro* metabolic stability assessment (*Supporting Information*) was performed by incubating compound **YF2** at a concentration of 1μM for 60 minutes in liver microsomes, with the aim to better understand the phase I metabolism in both species, as reported in **Table 2**. Compound **YF2** showed a poor half-life of 10 min in mice microsomes and 4.35 min in human microsomes with Cl_int_ of 0.137 ml/min/mg and Cl_int_ of 0.299 ml/min/mg, respectively. These values indicate that compound **YF2** is rapidly metabolized.

**Table 2.** In vitro metabolic stability of compound YF2 in mouse and human liver microsomes.

| Compound | Species | Half-life (min) | Clint (mL/min/mg protein) |
| --- | --- | --- | --- |
| YF2 | Human | 4.6 | 0.299 |
|  | Mouse | 10 | 0.137 |

In this context, to better understand the poor *in vitro* half-life, we investigated the metabolic profile in human microsomes on compound **YF2**, as reported in **Table 3**. The primary metabolic pathway for compound **YF2** is a demethylation reaction at the *N,N*-dimethylaminoethyl side chain, which accounts for 86.60% of the compound after 15.3 minutes of incubation in human microsomes. Additionally, 10.89% and 1.35% of compound **YF2** undergoes two hydroxylation reactions occurring on either phenyl rings. These data suggest that compound **YF2** is susceptible to phase I metabolic reactions, corroborating its poor *in vitro* microsomal stability.

**Table 3.** *In vitro* detection of metabolites of compound YF2 in human microsomes.

| Analyte (Biotransformation) | Retention Time, min | [M+H]<sup>+</sup>, m/z | Relative Level (% to Remaining Test Article) |
| --- | --- | --- | --- |
| **YF2** | 15.4 | 431.1331 | 100.00 |
| Demethylation | 15.3 | 417.1174 | 86.60 |
| Hydroxylation | 15.7 | 447.1280 | 10.89 |
| Hydroxylation | 13.8 | 447.1280 | 1.35 |
| Desaturation | 15.4 | 429.1174 | 0.81 |
| Hydroxylation + Demethylation | 18.4 | 433.1124 | 0.70 |

## 3. Discussion

In this study, we report the design, synthesis, and characterization of the activity of small molecules that modulate enzymatic histone acetylation. We also report the molecular docking and metabolic profile of one of these small molecules, **YF2**. Despite the common *N*-phenylbenzamide scaffold and the fact that the starting compounds CTPB and CTB were activators of p300, we discovered that changes in the side chain R^1^ could modulate the activity of the compounds towards opposite effects onto the substrate. Specifically, in contrast to **YF2, RA010168, RA010900**, and **RA010160** that behaved as p300 activators, compounds **JF1, JF10** and **JF16** inhibited the activity of the enzyme. A major difference between these compounds is related to the presence of the cyclic amine in **JF1** or longer and branched amino-alkyl and alkyl groups in **JF10** and **JF16**. Thus, we suggest that the size and structure of the alkyl group can significantly influence the biological activity of the compound.

Despite discovering both p300 activators and inhibitors, we selected **YF2** for further molecular modeling studies and metabolic profiling. Such a decision was motivated by the future directions of our investigation. For instance, enhancement of histone acetylation through HDAC inhibition has been shown to be beneficial in animal models of Alzheimer’s Disease (27). Indeed, the primary strategy used to up-regulate histone acetylation involves HDAC inhibition. However, the pleiotropic effect of nonspecific HDAC inhibition, combined with the opposite effects of HDAC 1 and 2 inhibition (28–31), may hamper the therapeutic potential of HDACIs. Therefore, we have focused on HATs as a target to up-regulate histone acetylation.

An initial literature analysis revealed the presence of two scaffolds for HAT activators. The first scaffold includes CTPB, its derivatives CTB and TTK21 (20, 21). The second one includes only one compound, nemorosone (4). CTPB and CTB activated p300 with low potency. Moreover, CTPB and CTB do not have favorable therapeutic characteristics as they are insoluble and membrane impermeable. Similarly, TTK21, which has a very similar structure to CTPB and CTB, must be conjugated to glucose-based carbon nanospheres (CSP) to cross the blood-brain barrier after intraperitoneal administration in mice (32). Nemorosone, a polyisoprenylated benzophenone derivative (PBD), is the only molecule among PBDs that enhances the p300 activity, exhibiting an 80% increase at 10 μM.

In summary, a new series of *N*-phenylbenzamide analogs were synthesized and evaluated for p300 enzymatic activity, docking properties, and metabolic profile. We identified p300 activators and inhibitors, using an *in vitro* screening approach. Among the *N*-phenylbenzamide derivatives, compound **YF2** exhibited a strong p300 activation profile but a limited *in vitro* microsomal stability. Nonetheless, compound **YF2** emerges as a lead in the pursuit of HAT activators. The discovery of compound **YF2** encourages us to pursue the drug optimization of this class of molecules.

## Acknowledgments

This work was financially supported by Appia Pharmaceuticals, the Kauffman (Ewing Marion) Foundation, the Alzheimer’s Association, and The Edward N. & Della L. Thome Award. The authors thank Appia Pharmaceuticals for providing compounds RA010160, RA010168 and RA010900, as well as Christopher Yarnold, Richard Jones, and Julian Rowley for designing them. All compounds in the RA series are proprietary to Appia Pharmaceutical and are protected by WO/2020/163731. The remaining compounds are protected through WO/2011/072243, WO/2012/088420, and WO/2015/153410 and have been licensed by Columbia University to Appia Pharmaceuticals. The authors thank Yuxuan Liu for helpful discussion on the method to measure enzymatic activity.

## Supporting Information

Supporting data include materials and methods, general procedures for intermediates and final compounds, and 1H and 13C NMR spectra.

## Author Contributions

JF, YF and RP designed compounds JF1, JF10, JF16, YF2 and RP102; EZ and JF performed the synthesis of the compounds used for the experiments described in the manuscript and wrote the manuscript; MM performed the molecular docking studies; EC established the method to measure enzymatic activity; EC, AS, BC and HZ performed the biochemical studies; EA,YIF, MF, EKA, CDB and HZ provided helpful discussion; SXD, DWL and OA supervised the work.

## Notes

### Competing Interest Statement

The authors have declared no competing interest.

