## Supporting Information for "Design and Synthesis of Potent First-in-Class Histones Acetyltransferase Modulators"

### 1. Materials and Methods.

All solvents and chemicals were reagent grade. Unless otherwise mentioned, all reagents and solvents were purchased from commercial vendors and used as received. Silica gel chromatography was performed using glass columns packed with silica gel (200−400 mesh, Aldrich Chemical). $^{1}H$ and $^{13}C$ NMR spectra were recorded on an Agilent-NMR-vnmrs 400 (400 and 100MHz, respectively) spectrometer and were determined in chloroform-d and DMSO-d6, with tetramethylsilane (TMS) (0.00 ppm) or solvent peaks as the internal reference. Chemical shifts (δ) are reported in ppm relative to the reference signal, and coupling constant values are reported in Hertz (Hz), multiplicity (s =singlet, d = doublet, dd = double doublet, t = triplet, q = quartet, m =multiplet, br = broad). The purity of all final compounds was ≥95% and was determined by LC/MS at wavelengths 220 and 254 nm. Liquid chromatography−mass spectrometry (LC/MS) was performed on two different instruments, a Shimadzu 2010A and a Shimadzu 2020 UFLC mass spectrometer, using a Waters Sunfire column (C18, 5 µm, 2.1 × 50 mm, a linear gradient consisting of 5-to-100% B over 15 min, followed by 100% B for 2 min, and then 5% B for 3 min (A = $H_2O$ (0.1% formic acid), B = $CH_3CN$ (0.1% formic acid)), flow rate 0.2 mL/min). HRMS (Electrospray Ionization) was performed at Columbia University, Department of Chemistry. Thin-layer chromatography (TLC) was performed on EMD pre-coated silica gel 60 F254 plates, and spots were visualized with UV light (254 nm). All reagents and solvents were used as received from major commercial suppliers, such as Sigma-Aldrich, Fisher Scientific, and Alfa Aesar without further purification. All air- or moisture-sensitive reactions were run under an atmosphere of argon in oven-dried glassware unless otherwise noted.

#### 1.1 *In vitro* Acetylation Assay on K18 and K27 on histone (H3).

To evaluate the biological activity of our library of compounds towards p300 enzyme, we developed an in *vitro* assay that was able to detect acetylated H3 compared to total H3 in the reaction mixture containing p300, H3, Ac-CoA, and compound. This assay allowed us to determine whether a compound was an activator or an inhibitor. A stock solution was prepared for each compound in either DMSO or $H_2O$, and a dilution concentration curve was generated. Reactions were carried out for 30 minutes in a buffer containing Tris-HCl, DTT, sodium butyrate, with the reaction initiated by adding the p300 enzyme. After 30 minutes, a mixture of coenzyme Ac-CoA and Histone substrate was introduced and incubated for an additional hour. Following the incubation, the samples were analyzed by SDS-PAGE and immunoblotting.

**Human Liver Microsomal Stability Assay.**

Mixed-gender human liver microsomes (Lot# 1010420) were purchased from XenoTech. The reaction mixture, minus NADPH, was prepared as described below. The test article was added into the reaction mixture at a final concentration of 1 μM. The control compound, testosterone, was run simultaneously with the test article in a separate reaction. An aliquot of the reaction mixture (without cofactor) was equilibrated in a shaking water bath at 37°C for 5 minutes. The reaction was initiated by the addition of the cofactor, and the mixture was incubated in a shaking water bath at 37°C. Aliquots (100 μL) were withdrawn at 0, 10, 20, 30, and 60 minutes. Test article and testosterone samples were immediately combined with 400 μL of ice-cold 40/60 acetonitrile (ACN)/H2O containing 0.1% formic acid and internal standard to terminate the reaction. The samples were then mixed and centrifuged to precipitate proteins. All samples were assayed by LC-MS/MS using electrospray ionization. The peak area response ratio (PARR) of analyte to internal standard at each time point was compared to the PARR at time 0 to determine the percent remaining at each time point. Half-lives and clearance were calculated using GraphPad software, fitting to a single-phase exponential decay equation.

**REACTION COMPOSITION**

- Liver Microsomes 0.5 mg/mL
- NADPH (cofactor) 1 mM
- Potassium Phosphate, pH 7.4 100 mM
- Magnesium Chloride 5 mM
- Test Article 1 µM

***In vitro* Detection of Expected Metabolites in Human Microsomes.**

Mixed-gender human liver microsomes (Lot# 1010420) were purchased from XenoTech. The reaction mixture, minus NADPH, was prepared as described below. The TA was added into the reaction mixture at a final concentration of 1µM. The control compound, testosterone, was run simultaneously with the TA in a separate reaction. An aliquot of the reaction mixture (without cofactor) was equilibrated in a shaking water bath at 37°C for 3 minutes. The reaction was initiated by the addition of the cofactor, and the mixture was incubated in a shaking water bath at 37°C. Aliquots (100 µL) were withdrawn at 0, 10, 20, 30, and 60 minutes. TA and testosterone samples were immediately combined with 400 µL of ice-cold 50/50 acetonitrile (MeCN)/$H_2O$ containing 0.1% formic acid and internal standard to terminate the reaction. The samples were then mixed and centrifuged to precipitate proteins. For metabolic stability determination, all samples were assayed by LC-MS/MS using electrospray ionization. For metabolite profiling, the T=60 minute incubation sample and the solvent control sample were analyzed using liquid chromatography (LC)-photodiode array (PDA)-high-resolution accurate mass spectrometry (HRAMS) (details of analytical method are outlined in Appendix 1). The peak area response ratio (PARR) to internal standard was compared to the PARR at time 0 to determine the percent remaining at each time point. Half-lives and clearance were calculated using GraphPad software, fitting to a single-phase exponential decay equation. Detection of expected metabolites, performed using MetWorks software (v. 1.3, Thermo), consisted of two steps: i) subtraction of the solvent control data file from the incubation sample data file using scaling factor 3 (generation of "SUB" file) and ii) automatic search and integration of the peaks in ion extracted chromatograms constructed using m/z values of the expected metabolites.

**General Procedure for Scheme of Synthesis.**

**5-hydroxy-2,2-dimethyl-4H-benzo[d][1,3]dioxin-4-one (2).** 2,6-dihydroxybenzoic acid (1) (10g) was dissolved in DME and stirred at 0°C. $SOCl_2$ was added dropwise and still at 0°C acetone was added dropwise. The reaction was stirred at room temperature for 24h. The reaction was quenched via addition of saturated solution of $NaHCO_3$. The mixture was poured into a separatory funnel, and extracted with ethyl acetate. The organic layer was separated and dried over $MgSO_4$. The solid was filtered off and the filtrate was concentrated under vacuum to afford a white precipitate, 98.3% yield. MS ESI (195.08m/z) $(M+1)^+$; $^1$H NMR δ (DMSO-$d_6$, 400 MHz): 10.29 (s, 1H), 7.51 (t, 1H, $J$ =7.99 Hz), 6.59 (dd, 2H, , $J_1$ =7.99 Hz, $J_2$ =3.99 Hz), 1.69 (s, 6H).

**5-alkoxy-2,2-dimethyl-4H-benzo[d][1,3]dioxin-4-one (3a-g).** 5-hydroxy-2,2-dimethyl-4H-benzo[d][1,3]dioxin-4-one (**2**) and $Cs_2CO_3$ were dissolved in 2-butanone, appropriate alkyl halide was added dropwise through a loader funnel. NaI was added at the mixture. The resulting mixture was refluxed for 24h and then the organic solvent was evaporated off. The crude was extracted with $CH_2Cl_2$ and NaOH and then washed with brine. The organic layer was separated, dried over $MgSO_4$ and evaporated under reduced pressure. The residue was treated with hot methanol to obtain a yellow pale precipitate. The desired compound was used in the following steps without any other purification. From 88.5% to 99.6% yield

**Alkoxy-6-hydroxybenzoic acid (4a-g).** 5-alkoxy-2,2-dimethyl-4H-benzo[d][1,3]dioxin-4-one (**3a-g**) was dissolved in THF, Cs(OH) was poured into the solution. The mixture reaction was stirred at reflux for 72h. The reaction was cooled down and diluted with ethyl acetate and then treated with HCl 1 N until pH=2. The white precipitate was filtrated, washed 3 times with $H_2O$ and dried under vacuum. The desired compound was used in the following step without any other purification. From 86.2% to 99.5% yield.

**General procedure amidation reaction (5a-g).**

Alkoxy-6-hydroxybenzoic acid (**4a-g**) and the appropriate aniline were dissolved in $CH_2Cl_2$ anhydrous at room temperature under argon atmosphere. EDC was added. The reaction mixture was stirred for 24h at room temperature under argon atmosphere. The solvent was evaporated off and the crude was treated with hot methanol and a white precipitate was obtained. The precipitate was washed with hot methanol and dried under vacuum to afford the desired product. Form 54.3% to 89.5% yield.

**2-ethoxy-6-hydroxy-N-(4-(trifluoromethyl)phenyl)benzamide (RA010160).**

White powder, 66.4% yield, LC/MS purity 98.1%, $t_R$= 6.32min; MS ESI (326.5904m/z) $(M+1)^+$; $^1H$ NMR δ ($CDCl_3$-d, 400 MHz): 13.42 (s,1H), 10.66 (s,1H), 7.92 (br s,1H), 7.75 (d, 1H, *J* =7.99 Hz), 7.49 (t, 1H, *J* =7.99 Hz), 7.41 (d, 1H, *J* =7.99 Hz), 7.32 (t, 1H, *J* =7.99 Hz), 6.68 (d, 1H, *J* =7.99 Hz), 6.45 (d, 1H, *J* =7.99 Hz), 4.28 (q, 2H, *J* =7.99 Hz), 1.65 (t, 3H, *J* =7.99 Hz). $^{13}C$ NMR

($CDCl_3$-d, 100 MHz): 169.02, 165.03, 157.78, 138.20, 134.23, 129.77, 123.88 ($^1J_{C,F}$ = 252 Hz), 121.37 ($^2J_{C,F}$ = 4.02 Hz), 117.60 ($^3J_{C,F}$ = 5.02 Hz), 112.38, 104.08, 102.22, 65.65, 15.02.

**N-(4-chloro-3-(trifluoromethyl)phenyl)-2-(2-(dimethylamino)ethoxy)-6-ethoxybenzamide (YF2).** Off white powder, 72% yield; LC/MS purity 99.8%, $t_R$=8.54 min; MS ESI (431.12m/z) $(M+1)^+$; $^1$H NMR δ (DMSO-$d_6$, 400 MHz): 10.76 (s,1H), 8.33(s, 1H), 7.91 (dd, 1H, $J_1$ =7.99 Hz, $J_2$ =3.99 Hz), 7.68 (d, 1H, $J$ =7.99 Hz), 7.38 (t, 1H, $J$ =7.99 Hz), 6.78(td, 2H, $J_1$ =7.99 Hz, $J_2$ =3.99 Hz), 4.41 (t, 2H, $J$ =3.99 Hz), 4.07 (q, 2H, $J$ =7.99 Hz), 3.41 (d, 2H, $J$ =3.99 Hz), 2.71 (d, 6H, $J$ =3.99 Hz), 1.23 (t, 3H, $J$ =7.99 Hz). $^{13}$C NMR (DMSO-$d_6$, 100 MHz): 163.65, 156.16, 154.98, 138.79, 132.17, 130.95, 126.45 ($^1J_{C,F}$ = 284 Hz), 123.86 ($^2J_{C,F}$ = 35.8 Hz), 117.44 ($^3J_{C,F}$ = 5 Hz), 106.16, 104.95, 64.04, 63.58, 55.27, 40.75, 14.54.

**2-(2-(dimethylamino)ethoxy)-6-ethoxy-N-(4-(trifluoromethyl)phenyl)benzamide (RA010168).** White powder, 55.4% yield; LC/MS purity 98.4%, $t_R$= 8.22min; MS ESI (397.1694m/z) $(M+1)^+$; $^1$H NMR δ ($CDCl_3$-d, 400 MHz): 11.81(br s, 1H), 9.52 (s, 1H), 8.26 (s, 1H), 7.97 (d, 1H, , $J$ =7.99 Hz), 7.43 (t, 1H, $J$ =7.99 Hz), 7.34 (d, 1H, $J$ =7.99 Hz), 7.24 (t, 1H, $J$ =7.99 Hz), 6.59 (d, 1H, $J$ =7.99 Hz), 6.50 (d, 1H, $J$ =7.99 Hz), 4.49 (br s, 2H), 4.04 (q, 2H, $J$ =7.99 Hz), 3.44 (br s, 2H), 2.78 (s, 6H), 1.34 (t, 3H, $J$ =7.99 Hz). $^{13}$C NMR ($CDCl_3$-d, 100 MHz): 207.20, 164.47, 157.04, 155.06, 139.51, 130.53 ($^1J_{C,F}$ = 178 Hz), 125.42, 122.90 ($^2J_{C,F}$ = 19 Hz), 120.44 ($^3J_{C,F}$ = 4 Hz), 116.36 ($^4J_{C,F}$ = 16 Hz), 106.38, 104.41, 64.65, 63.32, 57.26, 44.41, 14.72.

**N-(4-chloro-3-(trifluoromethyl)phenyl)-2-ethoxy-6-(2-(piperidin-1-yl)ethoxy)benzamide (JF1).** White powder, 67.4% yield; LC/MS purity 99.93%, $t_R$= 9.31min; MS ESI (471.15m/z) $(M+1)^+$; $^1$H NMR δ (DMSO-$d_6$, 400 MHz): 10.79 (s, 1H), 10.47 (br s, 1H), 8.34 (d, 1H, $J$ =3.99 Hz), 7.93 (dd, 1H, $J_1$ =7.99 Hz, $J_2$ =3.99 Hz), 7.70 (d, 1H, $J$ =7.99 Hz) 7.38 (t, 1H, , $J$ =7.99 Hz), 6.78 (dd, 2H, $J_1$ =7.99 Hz, $J_2$ =3.99 Hz), 4.43 (d, 2H, , $J$ =3.99 Hz), 4.07 (q, 2H, , $J$ =7.99 Hz),

3.38 (br s, 2H), 2.84 (q, 2H, , $J$ =7.99 Hz), 1.70-1.52 (m, 6H), 1.18 (t, 3H, $J$ =7.99 Hz). $^{13}$C NMR (DMSO-$d_6$, 100 MHz): 163.99, 156.11, 155.01, 138.81, 132.24, 131.00, 126.76 ($^{1}J_{C,F}$ = 282 Hz), 123.80 ($^{2}J_{C,F}$ = 27 Hz), 117.42 ($^{3}J_{C,F}$ = 7 Hz), 116.58, 106.16, 104.96, 64.07, 63.43, 54.75, 52.56, 22.37, 20.94, 14.58.

**N-(4-chloro-3-(trifluoromethyl)phenyl)-2-(3-(diethylamino)propoxy)-6-ethoxybenzamide (JF10).** White powder, 55.6% yield; LC/MS purity 96.3%, $t_R$= 9.42min; MS ESI (459.1817m/z) $(M+1)^+$; $^{1}$H NMR δ ($CDCl_3$-d, 300MHz): 7.94-7.82 (m, 3H), 7.46 (d, 1H, $J$=8.7 Hz), 7.31-7.25 (m, 1H), 6.57 (d, 2H, $J$=8.4Hz), 4.07 (t, 4H, $J$=6.6 Hz), 2.61 (t, 2H, $J$=7.2 Hz), 2.53 (q, 2H, $J$=6.9 Hz), 1.97-1.89 (m, 2H), 1.37 (t, 3H, $J$=6.9 Hz), 0.98 (t, 6H, $J$= 7.2 Hz). $^{13}$C NMR (DMSO-$d_6$, 100 MHz): 164.07, 156.26, 156.02, 139.07, 132.07, 130.73, 126.70 ($^{1}J_{C,F}$ = 299Hz), 123.57 70 ($^{2}J_{C,F}$ = 50Hz), 117.39 70 ($^{3}J_{C,F}$ = 6Hz), 116.76, 105.31, 105.09, 66.32, 63.93, 48.56, 46.25, 21.83, 14.61.

**N-(4-chloro-3-(trifluoromethyl)phenyl)-2-ethoxy-6-(isopentyloxy)benzamide (JF16).** White powder, 55% yield; LC/MS purity 99.64%, $t_R$= 16.567min; MS ESI (430.20m/z) $(M+1)^+$; $^{1}$H NMR δ (DMSO-$d_6$, 400 MHz): 10.57 (s, 1H), 8.25 (s, 1H), 7.91 (d, 1H,, $J$=7.99Hz), 7.66 (d, 1H, $J$=7.99Hz), 7.32 (t, 1H,, $J$=7.99Hz), ), 6.71 (dd, 2H, $J_1$=3.99Hz, $J_2$=7.99Hz), 4.07-3.98 (m, 4H), 169-1.62 (m, 1H), 1.48 (q, 2H, $J$=7.99Hz), 1.22 (t, 3H, $J$=7.99Hz), 0.78 (d, 6H, $J$=3.99Hz). $^{13}$C NMR (DMSO-$d_6$, 100 MHz): 164.06, 156.39, 156.04, 138.99, 132.06, 130.69, 126.66 ($^{1}J_{C,F}$=30.16Hz), 123.64 ($^{2}J_{C,F}$=8.04Hz), 117.50 ($^{3}J_{C,F}$=5.02Hz), 105.27, 105.22, 66.66, 63.92, 37.35, 24.40, 22.25, 14.59.

**N-(4-chloro-3-(trifluoromethyl)phenyl)-2-(2-(dimethylamino)ethoxy)-6-methoxybenzamide (RA010900).** White powder, 59.7% yield; LC/MS purity 97.8%, $t_R$= 8.40min; MS ESI (417.1148m/z) $(M+1)^+$; $^{1}$H NMR δ (DMSO-$d_6$, 400 MHz): 10.76 (s,1H), 8.33(s, 1H), 7.91 (dd, 1H, $J_1$ =7.99 Hz, $J_2$ =3.99 Hz), 7.68 (d, 1H, $J$ =7.99 Hz), 7.38 (t, 1H, $J$ =7.99 Hz), 6.78(td, 2H, $J_1$

=7.99 Hz, $J_2$ =3.99 Hz), 4.41 (t, 2H, $J$ =3.99 Hz), 4.07 (q, 2H, $J$ =7.99 Hz), 3.92 (s, 3H), 2.71 (d, 6H, $J$ =3.99 Hz). $^{13}$C NMR (DMSO-$d_6$, 100 MHz): 163.65, 156.16, 154.98, 138.79, 132.17, 130.95, 126.45 ($^1J_{C,F}$ = 284 Hz), 123.86 ($^2J_{C,F}$ = 35.8 Hz), 117.44 ($^3J_{C,F}$ = 5 Hz), 106.16, 104.95, , 63.58, 55.27, 40.75.
